# Comparative analysis of hsp16 and hsp70 heat shock protein families in *Caenorhabditis* nematodes

**DOI:** 10.1101/2025.10.13.682214

**Authors:** Wouter van den Berg, Harvir Bhullar, Bhagwati P. Gupta

## Abstract

The heat shock response (HSR), driven by molecular chaperons, is a key defense mechanism against proteotoxic stress. In nematodes, the hsp16 family and hsp70 family play central roles in the HSR, but their evolutionary conservation and expression patterns remain poorly understood. Here, we performed phylogenetic, genomic, and transcriptomic analyses of hsp16 and hsp70 genes in *Caenorhabditis elegans* and *C. briggsae*, and other related nematodes. Our findings show that hsp16 genes are rapidly evolving and often organized in clusters of closely spaced and oppositely oriented pairs. These genes show dynamic temporal expression in *C. elegans* and *C. briggsae* under basal, unstressed conditions. In contrast, hsp70 genes are more conserved across species and display stable expression. We also found that *C. briggsae* contains more orthologous HSP genes than *C. elegans*, and exhibits a higher proportion of constitutive expression, consistent with its greater thermal tolerance. These findings highlight the evolutionary diversification and functional organization of heat shock proteins in nematodes, offering insights into how genomic architecture and gene expression contribute to species-specific thermal adaptation.

## INTRODUCTION

Free-living nematodes can be found all over the world, and species have different environmental temperature ranges at which they are able to effectively reproduce and survive (Cutter 2015). The model organism *C. elegans* has a wide geographical range and is found in temperate and tropical climates (Andersen, et al. 2012; Felix and Duveau 2012). Laboratory studies have shown early on that the N2 Bristol reference strain, native to the U.K., is well-adapted to temperatures around 15-20**°C** but has a decreased lifespan and reproductive capacity at higher temperatures (Brenner 1974; Klass 1977). This temperature range appears to be a trait inherent to the species, as strains that were isolated from warmer climates do not significantly differ in their thermal tolerance (Petrella 2014). Other closely related nematode species such as *C. briggsae* and *C. inopinata* are better adapted to heat (Jhaveri, et al. 2025; Woodruff, et al. 2019).

An organism’s capacity to thrive at higher temperatures is dependent on how it deals with heat stress and the effect this has on protein stability. The HSR is a highly conserved guardian of proteostasis (Brehme, et al. 2014; Lazaro-Pena, et al. 2022). Heat shock proteins (HSP) are key components in the HSR, where they act as chaperones to preserve protein folding (Gu, et al. 2023; Leroux, et al. 1997). Like other higher eukaryotes, there are many heat shock proteins in *C. elegans* and other nematodes (www.wormbase.org). Foldase genes such as the hsp70 family contain two major domains: one that binds to unfolded proteins and the other that carries out folding and refolding in an ATP-dependent manner. Holdases, i.e. the small heat shock proteins (sHSPs) are thought to have evolved more recently (Huang, et al. 2008) and function by binding and preventing aggregation of unfolded proteins but lack the domain that facilitates refolding (Shim, et al. 2003; Veinger, et al. 1998). sHSPs co-aggregate, stabilize and prepare unstable and unfolded proteins for easier processing by hsp70 family member chaperones and other large HSPs (Goncalves, et al. 2021; Gu, et al. 2023).

sHSP and hsp70 genes are differentially involved in responses to a wide range of physiological and environmental stresses (Chen, et al. 2019; Hong, et al. 2004; Kuzmin, et al. 2004; Lin, et al. 2022; McElwee and Freedman 2011), but they are consistently involved in the HSR. Heat stress strongly induces expression of hsp16's, *hsp-17* and several hsp70 genes (Brunquell, et al. 2016; Candido 2002; Ezemaduka, et al. 2017; Hong, et al. 2004; Jovic, et al. 2017; Russnak, et al. 1983; Snutch, et al. 1988), and the genes *hsp-16*.*20*/*F08H9*.*3* and *hsp-16*.*21*/*F08H9*.*4* have also displayed an HSR, albeit to a lesser degree (Shim, et al. 2003).

Among the HSP genes, sHSPs of the hsp16 subfamily have undergone a significant expansion and show unique genomic arrangement. Many of the hsp16 genes are present as several pairs and are highly similar in sequence (Jones, et al. 1986; Shim, et al. 2003). Comparative genomic studies have been done with HSPs in other nematodes including *C. briggsae* (Aevermann and Waters 2008; Hong, et al. 2004). In comparisons between *C. elegans* HSPs and homologs in other nematodes, it remains unclear how conserved gene roles are, i.e. remaining constitutively active versus responding to various stresses. *C. briggsae* can survive and reproduce at comparatively higher temperatures than *C. elegans* (Felix and Duveau 2012; Poullet, et al. 2015; Prasad, et al. 2011). It also mounts a less intense HSR against mild heat shock (Jhaveri, et al. 2025). Furthermore, *C. briggsae* has a lower resistance to oxidative, osmotic and ER-UPR stress than does *C. elegans* (Jhaveri, et al. 2025), indicating that the two species have stress response systems attuned to different environments. This fits with previous findings that, geographically, the habitats of *C. elegans* and *C. briggsae* overlap but that relative population sizes are dependent on seasonal temperature changes (Felix and Duveau 2012). The transcriptomic background of these specialized adaptations remains to be elucidated.

Even under regular physiological conditions, in the absence of heightened levels of stress from heat or other sources, several members of the hsp70, hsp90 and sHSP gene families are constitutively expressed and have normal physiological functions (Agarraberes and Dice 2001; Kiefhaber, et al. 1991; Tang, et al. 2005; Welch and Feramisco 1985). In nematodes, *hsp-1* is an example of a constitutively expressed hsp70 member, whereas several others — *hsp-70, F11F1*.*1* and *F44E5*.*4* — are all expressed in response to stress (Brehme, et al. 2014; Nikolaidis and Nei 2004). Unlike hsp70 genes, little work has been done to investigate expression of hsp16 family members during standard, non-stressed conditions (Stringham, et al. 1992).

In this study we examined the conservation of HSP gene sequences and their genomic arrangements in nematodes. By performing phylogenetic analysis of the hsp16 and hsp70 gene families in six nematode species we show an evolutionary history of the heat shock gene proliferation. We show that sHSPs are prevalently clustered on single chromosomes in the studied nematode species. We also examined the longitudinal expression of hsp16 and hsp70 family members in *C. elegans* and *C. briggsae*, which collectively were revealed to have an overall higher rest-state gene expression in *C. briggsae*. Our findings suggest that hsp16 and hsp70 genes collectively contribute to higher thermal tolerance on *C. briggsae*.

## RESULTS

### Phylogeny of the hsp16 family in *Caenorhabditis* nematodes

We assembled an expanded *C. elegans* HSP gene set, consisting of hsp70 and sHSP genes, using InterPro domain annotations, literature on heat-shock chaperones, and BLAST matches (Brehme, et al. 2014; Brunquell, et al. 2016; Jovic, et al. 2017). The final list contains a total of 30 *C. elegans* HSP genes. To place these within an evolutionary context, we inferred orthology with OrthoFinder across five *Caenorhabditis* species (*C. elegans, C. briggsae, C. remanei, C. nigoni, C. inopinata*), another nematode *Pristionchus pacificus*, and the arthropod *Drosophila melanogaster. C. nigoni* and *C. inopinata* were chosen because these are sister species to *C. briggsae* and *C. elegans*, respectively, and *C. remanei, P. pacificus* and *D. melanogaster* were included as outgroups within the genus *Caenorhabditis*, within the phylum Nematoda and outside the phylum, respectively.

Sequence alignment and phylogenetic tree construction (Supplementary figure S1) resolved two hsp16 orthogroups that correspond to the *hsp-16*.*1*-like and *hsp-16*.*48*-like subfamilies (Fig. 1A; Supplementary Figure S1). *sip-1* and its orthologues are included in the hsp16 branch, but are in a distinct orthogroup, and in *C. elegans* expression is reduced under heat stress (Linder, et al. 1996). The heat-inducible sHSP *hsp-17* is more diverged from the hsp16’s, related by sequence yet falling on a separate branch. As expected based on species relationships, *C. briggsae* clusters with *C. nigoni*, typically with *C. remanei* as the next neighbor. *C. elegans* genes form cohesive groups within both hsp16 subfamilies, consistent with a most recent common ancestor (MRCA) of the *Caenorhabditis* genus carrying two hsp16 genes. No *D. melanogaster* genes are found on the hsp16 branch, suggesting hsp16 expanded after the Nematoda-Arthropoda split. The divergent *Drosophila* sHSP, *CG43851* (Morrow and Tanguay 2015), was not recovered in our analysis. In *P. pacificus*, 11 sHSPs cluster together and share the *hsp-16*.*48* orthogroup, indicating higher conservation of the *hsp-16*.*48*-like lineage relative to *hsp-16*.*1*-like genes. For context, hsp70 family relationships show deeper splits across *Caenorhabditis, P. pacificus*, and *Drosophila* (Fig. 1B), consistent with the older origin compared to the hsp16 family. Still, the *C. elegans* heat-responsive *hsp-70, F44E5*.*4* and *F44E5*.*5* form a *Caenorhabditis*-specific sub-branch that shares a single common ancestor with four *P. pacificus* orthologs further back, while the closest *Drosophila* orthologs are more closely related to the *hsp-1* lineage.

**Figure 1.**
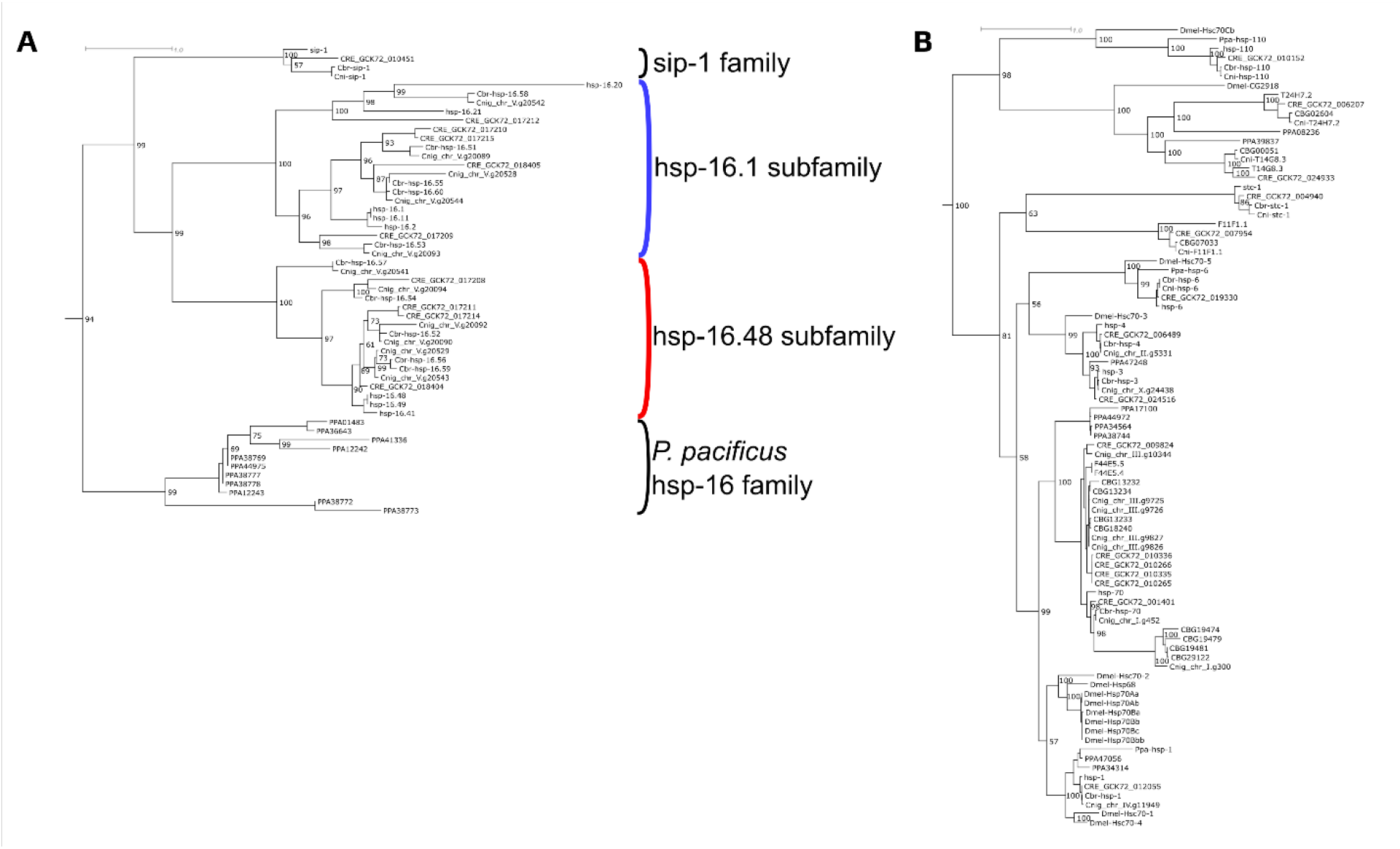
Phylogeny of hsp16 and hsp70 family genes among nematodes. Phylograms of **(A)** hsp16 gene family and **(B)** hsp70 gene family. Tree conformations are based on deduced amino acid sequences of all *C. elegans* hsp16 and hsp70 genes, as well as all homologs *from C. briggsae, C. remanei, C. nigoni, P. pacificus* and *D. melanogaster*. Trees were extracted from a larger tree based on all sHSP and hsp70 family genes, provided in Supplementary Figure S1. Colours indicate hsp-16 subfamilies in blue and red. Scale bar indicates branch length, i.e. nucleotide substitutions per site. Maximum likelihood was used to calculate parameters of the phylogenetic tree of sequence evolution, numbers adjacent to branches are bootstrap support values in percentages.

### Genomic organization of HSP genes

In *C. elegans*, the hsp16 genes are located on chromosome 5 in three distinct clusters (Figure 2A). The two clusters located on the left chromosome arm are made up of one and two pairs, respectively, of closely spaced genes with opposite orientations facing outwards. Cluster 1 contains the pair of *hsp-16*.*2* with *hsp-16*.*41*, and cluster 2 contains the pairs of *hsp-16*.*41* with *hsp-16*.*48* and *hsp-16*.*2* with *hsp-16*.*49*. The intergenic distances between the paired genes are short, between 200-400 bases. Previously it was shown that the intergenic regions between pairs are conserved and contain shared regulatory elements in *C. elegans* (Kay, et al. 1986; Stringham, et al. 1992) and in *C. briggsae* (Hong, et al. 2004). The expansion of the hsp16 gene family is likely due to the duplication of genes present in pairs. The third cluster consists of *hsp-16*.*20* and *hsp-16*.*21*, which are also closely spaced but not oppositely oriented.

**Figure 2.**
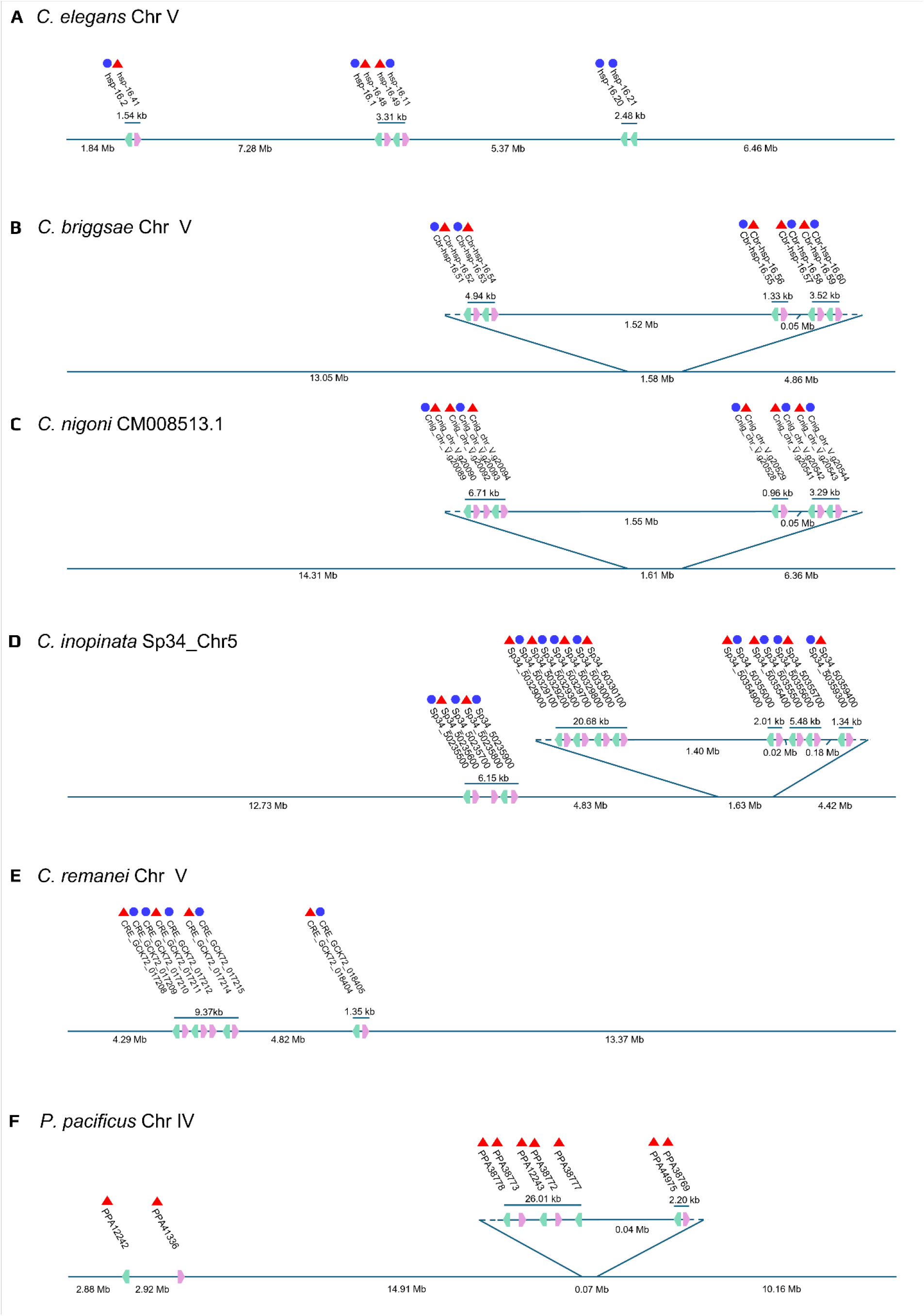
Chromosomal clustering of hsp16 family genes in *Caenorhabditis* species and *Pristionchus pacificus*. Chromosomal maps depicting relative locations of hsp16 family genes. Positions of genes are not drawn to scale. Genomic regions containing clusters of genes have been enlarged. **(A)** *C. elegans* Chromosome 5. **(B)** *C. briggsae* Chromosome 5. **(C)** *C. nigoni* Chromosome CM008513.1. **(D)** *C. inopinata* Chromosome 5 (Sp34_Chr5). **(E)** *C. remanei* Chromosome 5. **(F)** *P. pacificus* Chromosome 4. Hsp17 family genes, and suspected hsp17 family members *C. inopinata Sp34_50111700, C. remanei GCK72_018831* and *P. pacificus PPA01483* have been omitted. hsp16 family orthologs omitted on this overview are two clustered genes situated on *C. inopinata* Sp34_Chr1 and one gene on *P. pacificus* chromosome 3. Colours indicate *hsp-16*.*1*-like and *hsp-16*.*48*-like subfamilies in blue and red based on phylogenetic grouping and orthogrouping. All *P. pacificus* genes are marked as part of the *hsp-16*.*48*-like subfamily, because all members are part of the orthogroup of *hsp-16*.*48* and share a single phylogenetic branch and thus could not be divided into subfamilies. The direction of gene transcription is indicated by arrow direction and colour: pink indicates forward strand, green indicates reverse strand. Underscored values indicate the size of a gene cluster, measured as the minimal distance fully covering all genes within a cluster. Values below the line are intergenic distances, or distance to a chromosome end for the outermost values.

While in *C. elegans* hsp16 pairs are spread out over chromosome 5, in *C. briggsae* all ten hsp16 orthologues are more highly concentrated, to three clusters within a span of 1.58 Mb (Figure 2B). Within the clusters are five pairs of genes which consist of one gene each of the *hsp-16*.*1*-like and *hsp-16*.*48*-like subfamilies. Gene pairs are all oppositely oriented and have small intergenic distances of less than 200 bases. In *C. nigoni* the localization of hsp16's is highly similar to *C. briggsae* (Figure 2C). The only notable difference is an additional forward gene present in the first cluster. Based on sequence similarity, this gene, *Cnig_chr_V*.*g20092*, is likely to be a duplication of its neighbour *Cnig_chr_V*.*g20090*. Together these paralogues are homologous to *Cbr-hsp-16*.52. Beyond this, the conformations of hsp16 genes are identical between *C. briggsae* and *C. nigoni. C. inopinata* is the most closely related species known to *C. elegans* (Kanzaki, et al. 2018). Remarkably, its hsp16 family chromosomal organization is very unlike *C. elegans*, and more similar to *C. briggsae* and *C. nigoni* (Figure 2D). Uniquely, *C. inopinata* has a pair of hsp16 orthologs on its first chromosome. Furthermore, it has five clusters on Sp34_Chr5, containing 21 genes that include ten pairs of oppositely-oriented genes. Beside the first cluster, the remaining clusters are concentrated on a span of 1.63 Mb. Of the clustered genes, 11 are part of the *hsp-16*.*1* orthogroup. The remaining genes, including *Sp34_50111700* which is a likely candidate for an *hsp-17* homolog, belong to the *hsp-16*.*48* orthogroup. Among other species, *C. remanei* has two tight clusters, of 7 and 2 genes (Figure 2E). The two clusters are situated apart by 4.82 Mb, on the left arm of chromosome 5. The chromosome also features a lone gene, *GCK72_018831*, which can reasonably be designated as the *hsp-17* homolog based on the phylogenetic analysis. Lastly, *P. pacificus*, an outgroup to the *Caenorhabditis* genus, has a distinct hsp16 conformation. The chromosome features ten orthologs, among which two clusters of five and two genes can be identified on a small stretch of 70 kb (Figure 2F). Notably, all members in the first cluster are further apart than in most other pairs, with 4 to 6 kb between each. Three more orthologs are present at large intergenic intervals on chromosome 4 and one more on chromosome 3, which defies defining a clear homolog candidate to *hsp-17*.

In contrast to the large hsp16 gene clusters, hsp70 family members are broadly dispersed across the chromosomes (Supplementary File 1). In *C. elegans*, a notable tandem pair of oppositely oriented genes (*F44E5*.*4* and *F44E5*.*5*) exists on chromosome 2. Other hsp70 genes are far removed or isolated on separate chromosomes. Across related nematodes, hsp70 loci also exhibit species-specific mini clusters. *C. briggsae* has two clusters of three genes one of 9 kb on chromosome 1 with the same orientations and one of 8.4 kb on chromosome 3 with alternating orientations. *C. nigoni* and *C. remanei* each contain two back-to-back, oppositely oriented pairs on chromosome 3 of roughly 4.5 kb, with short intergenic regions (0.46 kb), alongside widely spaced loci on other chromosomes. *C. inopinata* harbors multi-gene clusters on chromosome 3 (19.3 kb) and chromosome 4 (13.7 kb) with mixed orientations, plus additional two genes within the next 0.25 Mb from the latter. In the outgroup *P. pacificus*, most hsp70 genes are dispersed, but chromosome 4 contains a small, 7.8 kb block of two genes.

Overall, hsp70 genes show a primarily dispersed distribution punctuated by short-spaced pairs and small clusters.

### Age-dependent expression of heat shock proteins in *C. elegans* and *C. briggsae*

Our longitudinal expression analysis of HSP genes revealed that under non-stressed conditions, hsp16 family transcripts in *C. elegans* fall into distinct age profiles (Fig. 3A). The central chromosome 5 cluster (*hsp-16*.*1, hsp-16*.*11, hsp-16*.*48, hsp-16*.*49*) did not display significant measurable expression across adulthood, whereas the adjacent pair *hsp-16*.*2* and *hsp-16*.*41* shows a shared, age-based expression pattern, increasing from day 1 to day 5 and then plateauing until day 10. The pair of *hsp-16*.*20* and *hsp-16*.*21* genes have dissimilar expression patterns, which may be due to their reverse orientation and, consequently, likely a lack of shared regulatory sequences. These data show that, in general, within-pair expression tends to be similar, while between-pair patterns show differences. Indeed, a previous study showed that this is also the case during heat shock (Stringham, et al. 1992).

**Figure 3.**
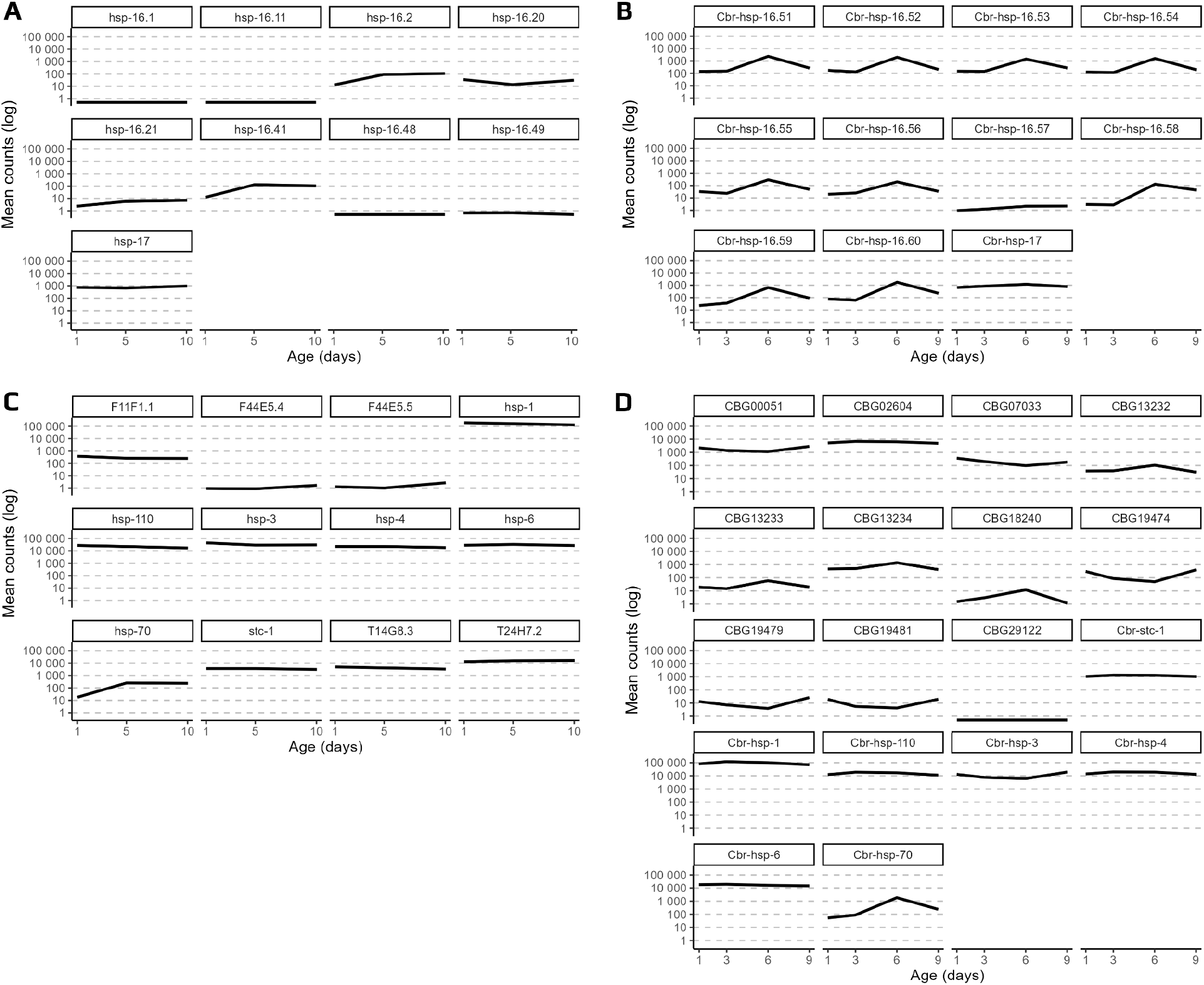
Expression of heat-stress responsive HSP gene families through nematode lifespan. Transcript counts of hsp16 and hsp17 genes in **(A)** *C. elegans* and **(B)** *C. briggsae*, and hsp70 genes in **(C)** *C. elegans*and **(D)** *C. briggsae*, on discrete timepoints from early to mid-life adult lifespan (X-axis). Lines are mean values of all replicates, formatted in log-scale (Y-axis).

In *C. briggsae*, expression values of the majority of hsp16's appear uniformly regulated and different from *C. elegans* genes, with expression increasing after day 3 but decreasing after day 6 (Fig. 3B). This matches a post-reproductive shift in stress-response programs (van den Berg and Gupta 2025). The only exceptions are *Cbr-hsp-16*.*57* which has consistently low expression and *Cbr-hsp-17* which maintains a uniform expression. *Cbr-hsp-16*.*57* and *C. elegans hsp-16*.*21* have similar basal profiles, but these are not shared by their adjacent gene pair partners.

Synteny and phylogeny of the two pairs of genes in *C. briggsae* and *C. elegans* do not cleanly resolve one-to-one relationships between these four genes. Thus, conservation of expression cannot be inferred without additional study of surrounding regulatory regions. By contrast, *hsp-17* orthologs in both species show stable, likely conserved basal expression, consistent with an early split of the *hsp-17* lineage from the rapidly diversified hsp-16 family.

For the hsp70 family, *C. elegans* genes can also be divided into several categories based on their constitutive expression levels (Figure 3C). Basal expression levels of *hsp-70* uniquely increase during the reproductive phase, whereas other hsp70 family members maintain stable levels of expression. Eight of the hsp70 genes (*hsp-1, hsp-3, hsp-4, hsp-6, hsp-110, T14G8*.*3, T24H7*.*2* and

*stc-1*) show consistently high levels of expression; *F11F1*.*1* is stably moderate; and the stress-responsive *F44E5*.*4* and *F44E5*.*5* remain negligible throughout. In *C. briggsae*, the one-to-one orthologs follow similar patterns to their *C. elegans* counterparts (Figure 3D). Similarly, highly expressed genes *CBG00051* and *CBG02604* are homologs to *T14G8*.*3* and *T24H7*.*2*, idem for *CBG07033* and *F11F1*.*1*. Importantly, *C. briggsae* carries eight orthologs related to *C. elegans F44E5*.*4* and *F44E5*.*5* and seven of these are expressed at varying levels with age, whereas CBG29122 is not expressed, highlighting expanded and diversified inducible modules in *C. briggsae*.

## DISCUSSION

Our comparative analyses across *Caenorhabditis* and outgroups clarify both the evolutionary history, genomic organization, and basal expression of heat shock protein families. Orthology and tree reconstruction revealed two hsp16 orthogroups (corresponding to *hsp-16*.*1*–like and *hsp-16*.*48*–like subfamilies) and six hsp70 orthogroups, with the hsp70 lineage distributed across deeper branches than hsp16. This arrangement supports a model in which hsp70 diversification predates the more recent expansion of hsp16 within *Caenorhabditis*. The hsp70 family branches have *P. pacificus* and *D. melanogaster* orthologs distributed among them, but the hsp16 family only has an isolated *P. pacificus* side branch and completely lacks representation in *D. melanogaster*. This arrangement is consistent with an older, more conserved hsp70 family and a younger, lineage-specific proliferation of hsp16 genes.

At the level of genome architecture, we find that hsp16 genes predominantly occur as oppositely oriented pairs with short intergenic distances (∼200–400 bp), arranged into species-specific clusters. *C. elegans* harbors three clusters on chromosome 5; *C. briggsae* and *C. nigoni* display denser clustering; *C. inopinata* extends this pattern with multiple clusters and an additional pair on chromosome 1; *P. pacificus* shows a distinct configuration with wider pair spacing in one cluster. These arrangements suggest an evolutionary history of local tandem duplications and a mechanism for co-regulation within pairs. By contrast, hsp70 genes show a more distributed chromosomal arrangement.

The expression datasets of hsp genes connect these phylogenetic and genomic arrangements to function. We found that under non-heat stressed conditions, most hsp16 genes in both *C. elegans* and *C. briggsae*, are constitutively expressed. In agreement with this, detectable expression of several family members was reported earlier in unstressed animals (Li, et al. 2019). Typically, these genes have been studied in heat-induced conditions and shown to exhibit high expression compared to non-stressed condition where their levels are either very low or below detectable limits (Baird, et al. 2014; Jones, et al. 1986). Thus, while hsp16 genes are involved in protecting animals against heat stress, we conclude that their basal expression also has biological function that is likely subject to additional regulatory inputs. We also found that hsp16 gene pairs in *C. elegans* frequently show similar expression trends (e.g., *hsp-16*.*2*/*hsp-16*.*41*), with some pairs showing high transcript counts and others remaining low throughout. A similar trend is observed in *C. briggsae* with corresponding age-dependent dynamics of hsp16 genes distinct from *C. elegans*. In both species, most hsp70 family members exhibit stable basal expression.

While the pair-centric expression dynamics of hsp16 genes is observed in both species, an earlier study reported that hypoxic stress caused differences in the expression of *hsp-16*.*1* and *hsp-16*.*41* pair as well as *hsp-16*.*2* and *hsp-16*.*48* pair (Hong, et al. 2004), providing evidence for additional unique roles of these genes outside of heat response. The functional significance of shared vs. distinct expression of hsp16 genes in response to different stress conditions remains to be investigated.

Taken together, the phylogenetic relationship, paired promoter architecture, and co-expression of paired genes lead us to suggest that duplicated and oppositely oriented pairs of hsp16 genes act as functional modules, while broader cluster position exerts less biological influence. This interpretation is consistent with prior reports of shared cis-elements in hsp16 intergenic regions and differential inducibility among pairs (e.g. see (Dixon, et al. 1990; Hong, et al. 2004). It also fits the observation that coordinated expression typically occurs within a pair rather than across an entire cluster. Studies have shown that genes with shared function, especially if co-expressed, often are located in close proximity on the genome (Beermann 1964; Roy, et al.

2002). There are several possibilities for this arrangement, including allowing the genes to share regulatory element binding regions or open chromatin state (Roy, et al. 2002; Tong, et al. 2017; Zhao, et al. 2019). A previous study showed that heat shock remodels chromatin at hsp16 loci and that open chromatin can allow the genes to transcribe more efficiently, and different sets of clustered genes display differences in these chromatin patterns (Dixon, et al. 1990). Our results are compatible with a scenario in which local promoter architecture is the dominant driver of basal co-expression, potentially augmented by shared open chromatin during induction. Distinguishing these mechanisms will require additional experiments such as targeted CRISPR edits of intergenic regions and cross-species promoter swaps.

Across our results, *C. briggsae* stands out with more *hsp* orthologs and a higher proportion of constitutive expression than *C. elegans*. These features are consistent with the greater thermotolerance exhibited by *C. briggsae* and suggest potential links between gene content, basal chaperone capacity, and heat resistance. The *C. briggsae* orthologs related to the stress-inducible *C. elegans* hsp70 genes (*F44E5*.*4*/*F44E5*.*5*) show age-dependent expression, implying that regulatory modules have diversified in ways that could tailor responses to the species’ ecological niche.

### Concluding remarks

Our data show that paired hsp16 modules expanded and diversified within *Caenorhabditis*, while hsp70 remained broadly conserved. The pair-centric architecture aligns with the basal co-expression of paired genes that we observe, and the broader constitutive chaperone capacity in

*C. briggsae* provides a plausible molecular basis for its higher thermotolerance. These data should prove valuable in future studies to understand how genomic organization and gene regulation shape species-specific responses to heat.

## METHODS

### Assignment of gene names

In consultation with Wormbase, the following HSP family members have been assigned gene names (Supplementary Table 1). These include *C. elegans hsp-16*.*20* (*F08H9*.*3*) and *hsp-16*.*21* (*F08H9*.*4*). The *C. briggsae* hsp16 genes have been named by order of occurrence on chromosome 5, as *Cbr-hsp-16*.*50 to 16*.*60* (11 genes in total), in consultation with Wormbase.

### Gene set selection

The hsp70 and sHSP gene sets were selected based on the presence of ‘HSP70’ and ‘small Hsp’ Interpro domains, respectively, as previously catalogued (Brehme, et al. 2014). Inclusion of further genes was based on two criteria. The first criterium involved genes which become more highly expressed following heat shock and facilitate survival under heat stress which includes *hsp-16*.*1, hsp-16*.*2, hsp-16*.*11, hsp-16*.*41, hsp-16*.*48, hsp-16*.*49, hsp-17, hsp-70, F44E5*.*4* and *F44E5*.*5* (Brunquell, et al. 2016; Iburg, et al. 2020; Jovic, et al. 2017). Nucleotide BLAST (blastn) performed on the CDS of all hsp16 and hsp70 genes found matches of > 65% sequence identity matches for *hsp-16*.*20* (*F08H9*.*3*) & *hsp-16*.*21* (*F08H9*.*4*) and for *F44E5*.*4 & F44E5*.*5*, respectively. Matches with pseudogenes were excluded.

### Phylogenetic analysis

For orthology analysis, proteome FASTA files were obtained for all included species. All files for nematodes were obtained from WormBase release WS295: *C. brenneri* PRJNA20035, *C. briggsae* PRJNA10731, *C. elegans* PRJNA13758, *C. inopinata* PRJDB5687, *C. japonica* PJRNA12591, *C. nigoni* PRJNA384657, *C. remanei* PRJNA577507 and *P. pacificus* PRJNA12644. *D. melanogaster* BDGP6.46 was obtained from Flybase release FB2025_01. Orthogroups were determined through Orthofinder, and *C. elegans* paralogs and all homologs to the selected set of *C. elegans* HSP genes were extracted (Emms and Kelly 2015). Amino acid sequences were obtained from proteome FASTA files for all candidate genes through custom R scripts from Ensembl release 113 for *C. elegans* and *D. melanogaster* (Harrison, et al. 2024), and from Wormbase WS295 for *C. briggsae, C. nigoni, C. remanei* and *P. pacificus* (Sternberg, et al. 2024).

Multiple sequence alignment was performed on the list of 201 hsp16 and hsp70 genes from *C. elegans, C. briggsae, C. nigoni, C. remanei, P. pacificus* and *D. melanogaster* in MUSCLE and T-COFFEE through the EMBL-EBI webUI (Madeira, et al. 2024) and output in CLUSTALW format. Aligned sequences were manually inspected for accuracy. Phylogenetic maximum likelihood analysis was performed in IQ-TREE 2 with UFBoot and 3000 bootstraps, numstop set to 1500 (Minh, et al. 2013; Minh, et al. 2020a, b). Best-fit model VT+G4+F model was used, determined by IQ-TREE internal ModelFinder. Tree visualization as phylogram was performed in Dendroscope (Huson and Scornavacca 2012). Gene family annotation and diagram tree rendition were made in Inkscape (https://inkscape.org/). For visual clarity, the prefix CRE_ was added to *C. remanei* gene names. *C. briggsae Cbr-hsp-16*.*58* and *C. nigoni* homolog *Cnig_chr_V*.*g20542* were separately included based on the presence of a ‘small Hsp’ Interpro domain, and *CBG26756* was removed due to false homology.

### Chromosomal mapping

Locations of all hsp16 genes in *C. elegans*, and all homologs in *C. briggsae, C. nigoni, C. inopinata, C. remanei* and *P. pacificus* were based on chromosomal coordinate values obtained through WormBase ParaSite (https://parasite.wormbase.org). CDS coordinates were used as gene location coordinates to exclude UTRs. Genes were considered to be clustered if present on the same chromosome with distances between genes smaller than 10kb (∼0.05% of chromosome size in *C. elegans*). Edges of gene pairs or gene pair clusters were determined to be coordinates of the furthest-apart nucleotides of the outer-most genes in the set, whether on the forward or reverse strand. Size of gene pairs or clusters were determined by subtracting the lowest edge coordinate value from the highest edge coordinate value, and distances between gene pairs or gene clusters were determined by subtracting the highest edge coordinate value of one set from the lowest edge coordinate value of the next set. Gene inclusion was based on orthogrouping with *hsp-16*.*1* and *hsp-16*.*48*, where the latter included *P. pacificus genes*. Hsp17 family genes and suspected hsp17 family members *C. inopinata Sp34_50111700, C. remanei GCK72_018831* and *P. pacificus PPA01483* were included in orthogroups, but were excluded from our analysis to focus on the hsp16 family.

### Expression in *C. elegans* and *C. briggsae* with age

Expressed gene counts were obtained from *C. briggsae* and *C. elegans* time-course using RNA-STAR and *featureCounts* as previously described datasets (Schmeisser, et al. 2013; van den Berg and Gupta 2025). Gene counts were averaged between replicates for each timepoint (*C. briggsae*: day 1, 3, 6, and 9 of adulthood; *C. elegans*: day 1, 5, and 10 of adulthood) and plotted with ggplot using custom R scripts.

## SUPPLEMENTARY DATA

**Supplementary Figure S1.**
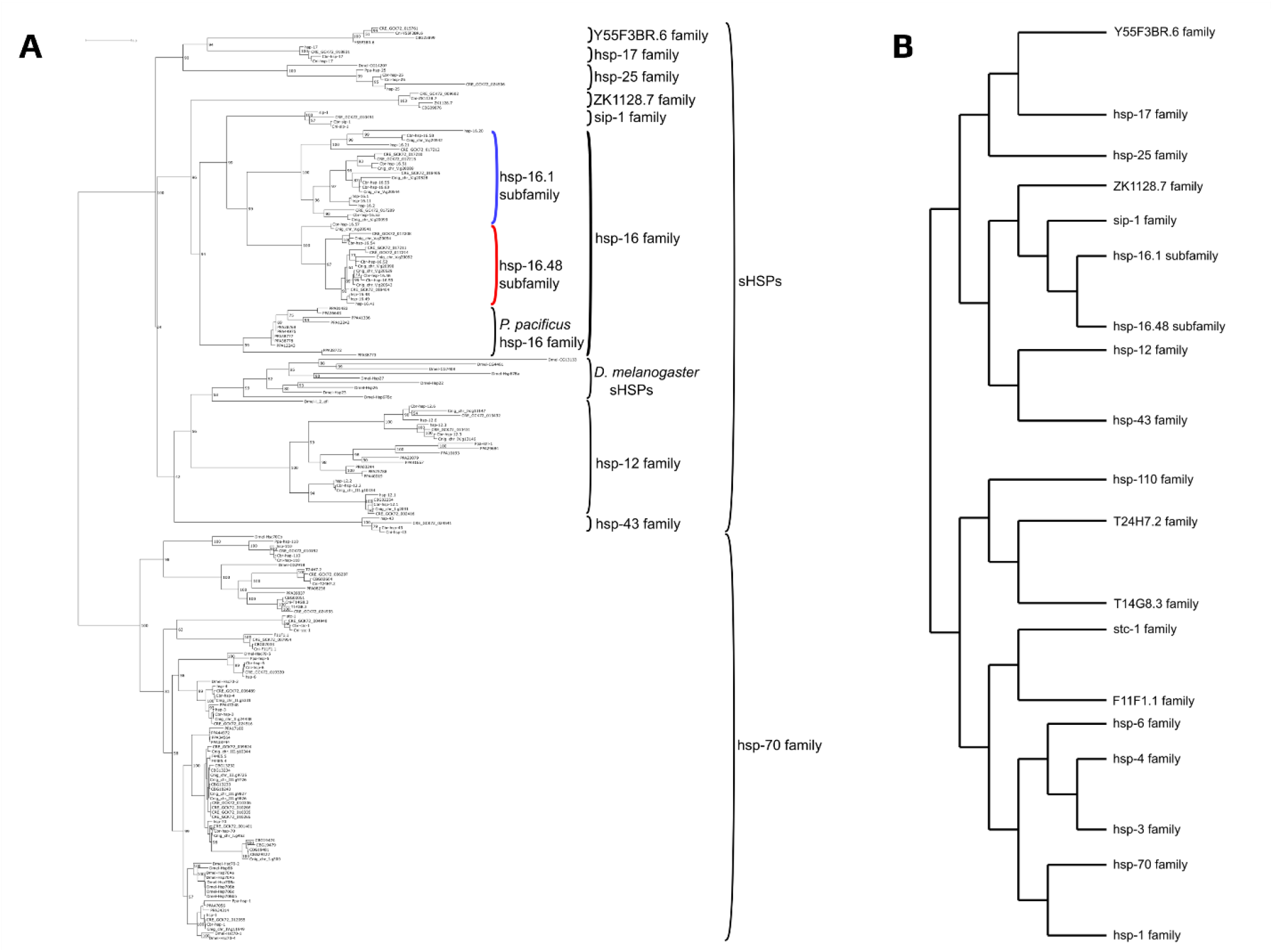
Phylogenetic relationships of sHSP and hsp70 family genes in nematodes. **A**. Phylogenetic tree of all sHSP and hsp70 genes in the nematodes *C. elegans, C. briggsae, C. remanei, C. nigoni, P. pacificus* and the fruit fly *D. melanogaster*. Gene family nomenclature is based on *C. elegans* genes and encompasses identified homologs in other species. Colours indicate hsp-16 subfamilies in blue and red. Scale bar indicates branch length, i.e. nucleotide substitutions per site. Maximum likelihood was used to calculate parameters of the phylogenetic tree of sequence evolution. Numbers adjacent to branches are bootstrap values in percentages. **B**. Diagram of sHSP & hsp70 phylogenetic tree to display *Caenorhabditis* gene families.

**Supplementary Table S1.** Public names of hsp16 family of genes corresponding to WBGene IDs and Sequence names.

| WBGene ID | Sequence name | Public name |
| --- | --- | --- |
| *C. elegans* |  |  |
| WBGene00008591 | *F08H9.3* | *hsp-16.20* |
| WBGene00008592 | *F08H9.4* | *hsp-16.21* |
| *C. briggsae* |  |  |
| WBGene00038447 | *CBG19184* | *Cbr-hsp-16.51* |
| WBGene00038448 | *CBG19185* | *Cbr-hsp-16.52* |
| WBGene00038449 | *CBG19186* | *Cbr-hsp-16.53* |
| WBGene00038450 | *CBG19187* | *Cbr-hsp-16.54* |
| WBGene00027233 | *CBG04591* | *Cbr-hsp-16.55* |
| WBGene00027234 | *CBG04592* | *Cbr-hsp-16.56* |
| WBGene00027244 | *CBG04605* | *Cbr-hsp-16.57* |
| WBGene00027245 | *CBG04606* | *Cbr-hsp-16.58* |
| WBGene00027246 | *CBG04607* | *Cbr-hsp-16.59* |
| WBGene00027247 | *CBG04608* | *Cbr-hsp-16.60* |

